# GeneSNAKE: a Python package for benchmarking and simulation of gene regulatory networks and perturbation-induced expression data

**DOI:** 10.1101/2025.11.10.685583

**Authors:** Thomas Hillerton, Anton Björk, Nils Lundqvist, Erik K. Zhivkoplias, Mateusz Garbulowski, Erik L. L. Sonnhammer

**Affiliations:** Department of Biochemistry and Biophysics, Stockholm University, Science for Life Laboratory, Box 1031, 17121 Solna, Sweden

**Keywords:** gene regulatory network, data simulation, ODE model, Hill equation, benchmarking, gene expression, gene perturbation, noise model

## Abstract

Understanding how genes interact with and regulate each other is a key challenge in systems biology. One of the primary methods to study this is through gene regulatory networks (GRNs). The field of GRN inference however faces many challenges, which necessitates effective tools for evaluating inference methods. For this purpose, data that corresponds to a known GRN, from various conditions and experimental setups is necessary, which is only possible to attain via simulation. Today, most existing tools for GRN-based simulation are limited either in network or data properties, with few or no options to modify these properties.

To address these limitations we present GeneSNAKE, a Python package designed to allow users to generate biologically realistic GRNs, and expression data for benchmarking purposes. GeneSNAKE allows the user to control a wide range of network and data properties, including several distinct noise models. GeneSNAKE improves on previous work in the field by adding a perturbation model and a wide range of perturbation schemes along with the ability to control the noise and the perturbation strength.

For benchmarking, GeneSNAKE offers a number of functions both for comparing network similarity, and properties in data and GRNs. These functions can further be used to study properties of biological data to produce simulated data with more realistic properties. GeneSNAKE is an open-source, comprehensive simulation and benchmarking package with powerful capabilities that are not combined in any other single package, and thanks to the Python implementation it can be extended and modified by users.

## Introduction

Understanding how genes interact with and regulate each other has long been one of the key challenges in systems biology. One of the primary methods to study this is through gene regulatory networks (GRN). The field of GRN inference however still faces many challenges, such as the complexity of gene regulation, high noise levels of expression data, and that a majority of the true regulatory interactions are unknown making any finding hard to verify. With these challenges in mind, a large number of tools have been developed to tackle the issue of gene regulatory network inference (GRNI), using a variety of statistical and machine learning approaches with a focus on different types of data and experimental approaches to capture the regulatory connections [1]. Therefore, a large number of methods, all with slightly different focus, necessitates effective tools for evaluating inference methods. For this purpose, data that corresponds to a known GRN and is derived from cells at similar conditions and cell cycle phase yet from various experimental conditions and setups would be required. A key issue with this approach arises here in the form of limited coverage of experimentally validated GRNs - for instance the human GRN in TRRUST is limited to 8,444 regulatory interactions for 800 transcription factors [2]. Due to this, obtaining data for a known GRN to evaluate GRNI methods is today impossible due to the excessive number of experiments that would be required both to properly determine the exact underlying GRN and obtaining informative data for meaningful comparison. This is often reflected in benchmarking papers where methods tested on real data often have a correctness so low that it can often not be distinguished from random performance [3] . Because of this the GRN field traditionally relies on synthetic data for developing and evaluating methods [4,5].

The need for synthetic data has led to a number of popular tools for gene expression simulation. The most popular option for simulators is the GeneNetWeaver (GNW) tool published by Shaffter et al in 2011 [6]. However, both earlier and later simulators [7–10] have been developed in an attempt to address the lack of data for GRNs. While these tools often perform well for the task at hand, they often suffer from being overly focused on a single issue of the data generation. For example, GNW focuses heavily on selecting a biologically realistic GRN to generate the data from. This however, comes at the expense of having very limited control over data properties and experimental design. The tool GeneSPIDER [7] allows the user to control the noise level and condition number when generating data, but can only simulate data at steady state. Another example is the BoolODE model developed by Pratapa et al [8] for simulating single-cell gene expression data given a GRN, but it does not have built-in functionality for the user to generate a GRN, and is limited to using boolean GRNs. While this specialization is not necessarily negative it does limit the applicability of the method as there is still no consensus in the GRN field of what properties are most important for GRNI. In the case of GNW, knockdown perturbations are limited to a fixed strength of 0.5, and for BoolODE no perturbation design can be specified. These shortcomings create a highly limited view when used for testing GRNI methods, as not all biological experiments follow a similar design or degree of perturbation. Finally, a recurring issue is that the simulator is hard to access either due to a lack of documentation, the code not being available, or the simulator being written in a language rarely used in bioinformatics. This is something that can either discourage or completely prevent the usage of a simulator for groups working with developing or studying GRNs.

Here we present GeneSNAKE (Generation and Simulation of Networks and datA pacKagE), a python package for generation of synthetic data that follows the dynamics specified in a GRN model. GeneSNAKE builds on previous works relying on a commonly used ODE model but offers further functionality aimed at improving GRN generation, experimental design, and dynamic user-defined data properties, see **Figure 1**. To address weaknesses of previous methods a focus has been placed on ensuring a balance between biological realism and flexibility, see **Table 1**. This has been ensured both by creating functions that generate GRNs and expression data that are biologically feasible and by allowing users to modify most parameters to control the generation in great detail. Further to allow for robust and varied experimental design when it comes to perturbing the system, GeneSNAKE offers a variety of predefined perturbation designs that can be used to generate data. GNW only allows perturbations at fixed strengths for knockdown (50%) and knockout (100%), respectively. For maximum flexibility, GeneSNAKE allows the user to choose perturbation strength per gene anywhere between 100% knockdown and infinite overexpression to simulate data with any experimental design. Finally, to offer as much flexibility as possible, GeneSNAKE includes multiple noise models that capture biologically relevant noise in the data and allow the users to customize the noise degree such that various experimental conditions can be approximated.

**Table 1.** Comparison of selected tools for simulating GRN-based gene expression data. ODE: Ordinary differential equations. SDE: Stochastic differential equations.

| Tool | GeneSNAKE | GeneSPIDER | GeneNetWeaver | BoolODE |
| --- | --- | --- | --- | --- |
| Authors | Hillerton et al. 2023 | Tjärnberg et al. 2017 [7] | Schaffter et al. 2011 [6] | Pratapa et al. 2020 [8] |
| Programming language | Python | Matlab | Java | Python |
| Distribution | Source code | Source code | Executable program | Source code |
| GRN generation | Internal with flexible topology and motif properties | Internal with limited topology properties | Loads external GRN | No |
| Data simulation | Dynamic ODE | Linear ODE approximation | Dynamic ODE | Dynamic ODE |
| Noise models | - Gaussian multiplicative<br>- Log-normal multiplicative<br>- Single cell sequencing like | - Gaussian additive | - Gaussian and log-normal additive.<br>- SDE model multiplicative<br>- microarray-like | - SDE model multiplicative<br>- dropout model |
| Perturbation strength and design | Tunable with customizable and pre-defined designs | Tunable with customizable design | Fixed with single or all-genes design | Stochastic with all-genes design |
| Maintained | yes | yes | no | yes |

**Figure 1.**
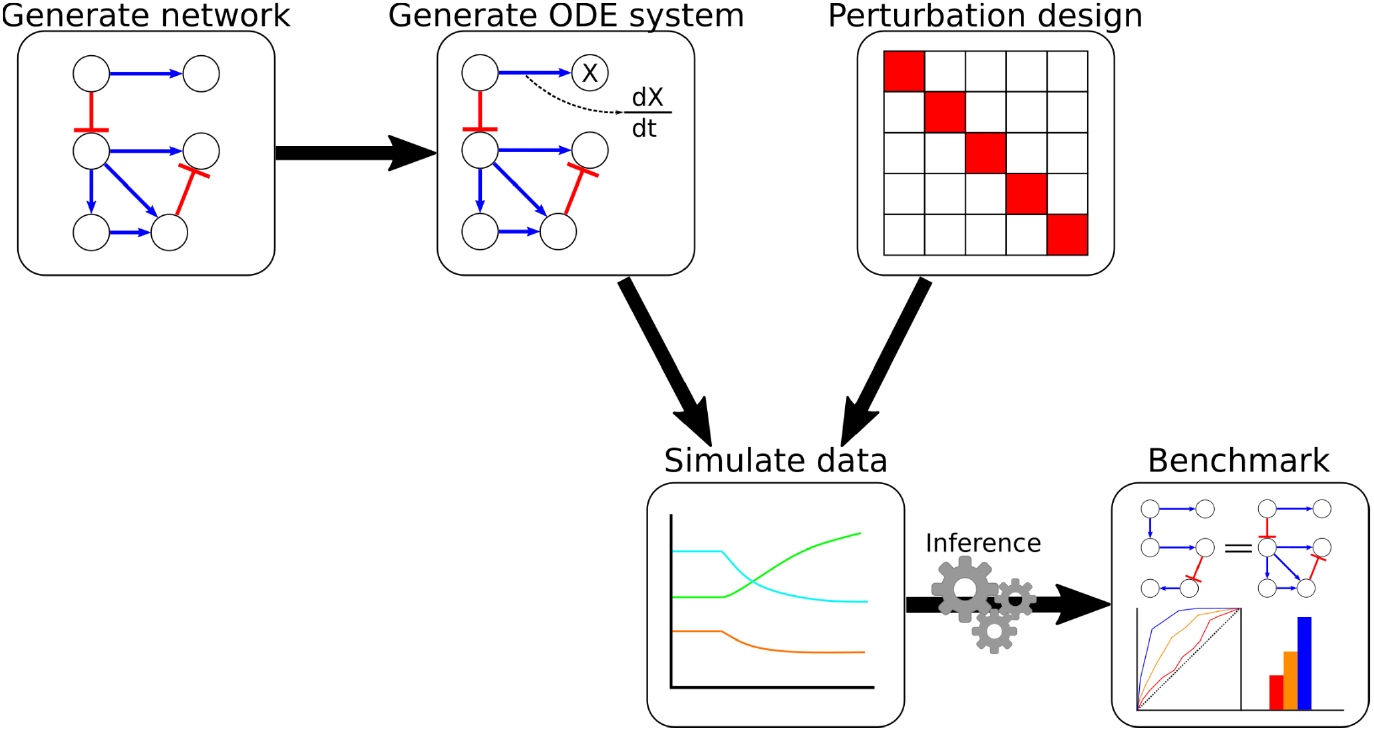
GeneSNAKE workflow. GeneSNAKE can generate GRNs, and from these, an ODE system that is used to generate time series and steady-state gene expression data following a large number of possible perturbation designs. The generated GRNs and simulated data feed into a benchmarking module that has a range of methods to compare GRNs inferred from the data with the original GRN.

## Methods

### Generating GRNs

GeneSNAKE simulates interactions between genes based on a GRN ensuring that any change in the system follows the dynamics described in the GRN. Thus it is important to have access to GRNs with different properties and sizes to simulate data from. Users can input any network to the algorithm in the form of an edge list or adjacency matrix. GeneSNAKE supports the generation of biologically realistic GRNs using the FFLatt algorithm that ensures a motif composition typical for biological GRNs [11]. It does this by employing a motif-based preferential attachment algorithm with a power-law kernel, aiming to reproduce the feed-forward loop (FFL) motif occurrence observed in literature-based biological GRNs [12]. It also preserves important topological properties such as scale-free topology, sparsity, and average in- and out-degree per node. The FFL-enriched network is generated with attachment rules and predetermined probabilities that control for the emergence of new FFL motifs at every iteration. The algorithm also allows for depletion of other three-node motifs, such as cascades, downlinks, uplinks, and cycles. GeneSNAKE further supports integration of any Networkx graph. Networkx is a commonly used graph handler in Python [13]. To obtain an overview of installation and how-to guidelines we encourage the user to visit the GeneSNAKE tutorial website: https://sonnhammer-tutorials.bitbucket.io/genesnake.html. The tutorial includes step-by-step examples, as well as descriptions of the GeneSNAKE parameters.

### Modeling gene expression as a molecular system

Model assumptions must be made on how the real system behaves and how closely the real system must be mimicked to make it useful. For GeneSNAKE several assumptions are made based on how genes are expressed and regulated. The most fundamental assumption made when creating the GeneSNAKE model is that gene expression works like a probabilistic molecular system that exists freely in solution. In practical terms, this means that gene expression is described as a sigmoid function, where activation and repression of a target gene is dependent on the concentration of the regulator gene, through a Hill equation where both minimum and maximum expression is bounded. The model assumes that transcription and translation occur in the same space, hence no molecular transport is required. This assumption is based on findings indicating that the time for translocation of mRNA and proteins is significantly lower than the half-life of molecules in the cytoplasm, <10 minutes and several hours respectively, in mice [14]. Further, the model assumes that the ribosomal concentration is always higher than the mRNA concentration and thus the relation between protein concentration and mRNA concentration for a given gene can be described by a linear differential equation where each mRNA molecule for a given gene is transcribed once per time step. This relationship ensures that the protein expression follows a similar sigmoid distribution as the mRNA, ensuring that the expression is bounded within the system. Finally, degradation of both protein and mRNA is modeled as a linear function based on a constant degradation rate for each molecule, with the assumption that the degradation systems for protein and mRNA always have more capacity than their maximum expression in the system. As degradation rates vary between different genes, we assign a random value to the degradation constant (λ) to make some genes degrade faster than others. The degradation rate for proteins is assigned to a lower value than for mRNA molecules on average to simulate the longer average half-life of proteins [15]. With these assumptions, gene expression can be modeled with the two-step ODE model developed for the DREAM3 challenge [16]. This model was selected as the basis for GeneSNAKE due to its frequent and well-established use in the GRN field. It is also employed by GeneNetWeaver [6] and BoolODE [8], two popular simulation tools in the GRNI field.

### Ordinary differential equations describing gene regulatory interactions

Using the above described assumptions on how genes are expressed and interact within a GRN it becomes possible to simulate the dynamic behavior in the system using two ordinary differential equations (equation 1 for mRNA and equation 2 for protein). The change in mRNA concentration (equation 1) for a given gene (*x*_*i*_) is described as the addition of new molecules, calculated as the max transcription rate (*m*_*i*_) times the protein concentration of each regulatory gene (*y*_*reg*_) given by the function *f(y*_*reg*_), minus the degradation of existing molecules, calculated as the degradation constant *λ*^*mRNA*^ times the current mRNA concentration *x*_*i*_.

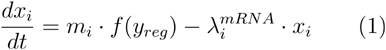

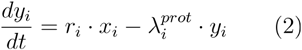

For the GeneSNAKE model, *m*_*i*_ is set to 1 for all unperturbed genes in the system and *λ*_*i*_^*mRNA*^ is randomly assigned a value from a uniform distribution between 0.2-0.7. The activation function *f(y*_*reg*_) depends on the protein concentration (*y*) of each regulatory gene and consists of a Hill function (equation 3). As the GeneSNAKE model assumes that all interactions occur through protein-mediated interactions, the change in protein concentration (y) over time is also modeled (equation 2). This extra step allows the model to capture the time lag between a change in mRNA concentration and the corresponding change in protein concentration.

The change in protein concentration (equation 2) is described by the number of new proteins translated, calculated as the max translation rate (*r*_*i*_) times the mRNA concentration (*x*_*i*_), minus the number of degraded protein molecules, calculated as the degradation rate 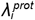 times the number of existing protein molecules. 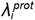 is randomly assigned to a value between 0.2-0.4.

To allow for complex interactions within the system, GeneSNAKE models regulations where 2 or more regulators interact, using an additive Hill function, shown for 2 regulators in equation 4. For the combinatory activation, the total activation is given by the effect from each regulator plus that of the combined, multiplicative effect of the regulators acting together. To model the combinatory effect, a combination factor *P* is used. In GeneSNAKE *P* is assigned to either 1 or 0, indicating either a combination that requires all participants, simulating an AND logic, or no combinatory effect, simulating an OR logic. In equation 5 this is done by a multiplication operator to get the AND logic, and addition to get OR. The combinatory effect is in GeneSNAKE limited such that all participants must be either activating or inhibiting, and no combinations can be made of regulators with opposing effects. For all of the activation/inhibition functions ( equation 3-5) GeneSNAKE uses the same parameters; α describes the regulatory effect in how strongly the target expression is affected, k is the dissociation constant, roughly equivalent to how likely it is that a regulator molecule does not have an effect, and n is the Hill coefficient, describing how quickly the regulator changes the expression. In GeneSNAKE these parameters are randomly assigned: k is set to a value between 0.2-0.6, and n to an int value between 2-4 (traditional range of values that will give a hill function). α is a special case being assigned values based on network properties. If the edge suggests a strong interaction (defined as an edge value greater than the median absolute edge weight in the network) then α is assigned a value between 0.6-1.0. If instead the edge is a weak interaction (edge value less than or equal to the median absolute edge weight) then α is assigned to a value between 0.4-0.8. Finally, if there is a collaborative interaction between two or more regulators (P=1) each regulator is assigned a weak α, i.e. 0.2-0.4, on their own and a strong α_3_, i.e. 0.7-1.0, for the combination.

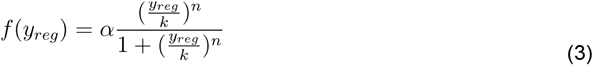

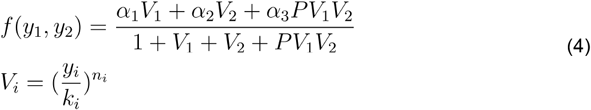

By enforcing a strict mRNA to protein to regulatory effect, the model can simulate dynamic time-delayed chains of reactions following the edges of the GRN, allowing for simulation of gene expression in both time series and steady-state experiments. The dynamic non-linear nature of the model allows for simulation of both complex and simple relationships between genes with multiple regulators collaborating or acting independently as desired by the user.

For ease of implementation and future-proofing, GeneSNAKE relies on the Python package Scipy [17] for solving ODE systems. Specifically, it uses the solve_ivp function from Scipy relying on LSODA, a Fortran-based ODE solver [18]. To ensure that the solution is robust the error tolerance was set to 1*10^-8^ to minimize errors in the simulation.

### Simulating perturbations

A key requirement for inferring GRNs from gene expression is the perturbation of the system to capture the system there must be a change that can be detected and explained. To simulate perturbations, GeneSNAKE modifies the max transcription rate parameter of equation 1 (m_i_) in the following way:

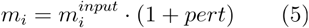

Where pert is a value between -1 and infinity, and m_i_^input^ is the original max transcription rate value used to obtain the control expression, by default 1. This models the perturbation by setting pert, the wanted change, to a non-zero value. For instance, pert = -0.8 gives an 80% reduction of the original transcription rate in the control. Several perturbation schemes have been proposed in the literature, but it is not clear which one is the best from a GRNI perspective. Because of this, GeneSNAKE features a wide variety of options for the generation of perturbation schemes as well as support for using custom perturbation designs. Currently, GeneSNAKE supports the perturbation schemes:

- Single gene - Each gene in the system is perturbed one by one so that all genes in the system are perturbed once.
- Combinatory - Multiple genes are perturbed in each experiment in such a way that each gene is perturbed roughly an equal amount of times across all experiments. The number of perturbed genes per experiment is determined by the user.
- Targeted combinations - Multiple genes are perturbed in each experiment and combinations are selected from the GRN so that genes regulating the same target are perturbed in the combinations. The number of perturbed genes per experiment is determined by the user.
- All genes - All genes in the system are perturbed with a random but small effect, either positive or negative, to simulate global perturbation experiments such as cell medium or temperature changes.
- Specific genes - A single or multiple genes are perturbed in each experiment based on an input list of gene names and gene combinations to perturb. Only those genes in the list will be perturbed.

For all of the perturbation schemes, excluding single perturbation, the number of perturbed genes per experiment is only limited by the total number of genes and the number of simultaneously perturbed genes. For targeted combinations, the algorithm will generate combinations with fewer genes in them if it is not possible to find combinations with the requested number of genes. This is to ensure that as many unique experiments as possible are created. It makes sure that each experiment has a unique perturbation scheme, except for when using the random perturbation scheme where experiments can be identical. The experiment parameters are of particular use for the combinatory and targeted perturbation scheme where possible combinations of genes will be selected at random until the maximum number of experiments have been achieved. Utilizing these schemes allows for high flexibility in the experimental design while automating its creation allowing for quick simulations of complex studies in a user-friendly manner. To make more elaborate experimental designs it is possible to merge multiple perturbation schemes and use this to simulate data, for example, to use a design of single gene perturbation of all genes along with a set of interesting combinations to be simulated.

### Noise modeling

Noise is one of the hardest obstacles for GRNI. Unfortunately, the exact nature of biological noise is today undetermined, and therefore GeneSNAKE offers several noise models as well as an option for the user to input either a noise function or a matrix of noise to add. Currently, three noise models are available in GeneSNAKE. The first one is a Gaussian noise model where the signal in each experiment is multiplied with noise drawn from a Gaussian distribution. Second, a log-normal noise model, previously shown to model microarray noise well [6],[19,20], where log-normal distributed noise is multiplied with the signal. Finally, a single-cell sequencing noise model based on a zero-inflated negative binomial noise distribution. This model uses a simulated coverage, gene length, error rate, and dropout rate to model biases in single-cell sequencing experiments (for details see Supplementary materials and Figure S2).

### Simulating an experiment

Using the above described ODE, perturbation scheme, and noise models GeneSNAKE simulates any number of experiments for measuring gene expression. Despite all the different options, the simulation of experiments is consistent, i.e. each simulation starts with the creation of a control expression of genes at steady-state. Steady-state here is determined by applying a multidimensional root-finding test ensuring that the change in the system has converged to a steady-state. GeneSNAKE then simulates each experiment in the perturbation design matrix by applying the selected perturbation to the gene expression and solving the ODE system using the control values as the starting point. The ODE system is then solved either for a given time span or until a new steady-state is detected by the Kalman filter in the system, depending on if the user requested generating time series data with multiple time points or steady-state data with only a single time point as output.

Once the simulation is complete, GeneSNAKE returns the experimental and control data along with the used noise for each generated time point. For time series data, to create biologically realistic simulations only a limited number of timepoints is returned, for which the exact number can be set by the user. The time points are selected either in a uniform linear distribution across the whole simulation time or in a log distribution with a diminishing number of timepoints as the time increases. For steady-state experiments, only a single time point is returned at the stopping point determined by a point where no gene or protein has moved more than a tolerance value for at least two time steps. It is also possible to use this as a stopping point for time series data to ensure that all the time points are located in the shifting part of the experiment rather than in steady-state areas. If no steady-state can be found due to a cyclic behavior in the system, steady-state data will instead be defined as the last time step in the model.

### GRN inference

GeneSNAKE comes with several methods for GRN inference, including LSCO [21], LSCON [22], LASSO [23,24], and Z-score [25]. Z-score is implemented in two variants, either using all genes (Z-score) or excluding the perturbation target gene from the estimation of gene expression mean and standard deviation (Z-score-P). This to avoid bias from the intentionally perturbed values, which are not representative of the observed distribution. In addition, GENIE3 [26] was used for investigating inference method performance against SNR levels in this publication, as an external Python package from https://github.com/vahuynh/GENIE3 and not included in GeneSNAKE. The publication reproduction code therefore includes separate setup of GENIE3. All methods were run with default parameters with a few exceptions. The *L*_*1*_ penalty in LASSO was set with Brent’s method to produce a sparsity of 50 links/gene on average. It has been shown that when running GENIE3 on perturbation-based data, the results are substantially better when the predicted GRNs are transposed, i.e. reversing the direction of regulator-target pairs, hence this was done here.

### Benchmarking

For GRN inference performance evaluation, GeneSNAKE computes and plots Receiver Operating Characteristic (ROC) curves, Precision Recall (PR) curves, and the Areas Under (AU) them, resulting in AUROC and AUPR values. For comparability across methods and situations, the curves are extended when predicted networks are incomplete. For ROC curves, the standard linear extension to (1, 1) is applied. For PR curves, the extension is done according to the DREAM 2 challenge performance evaluation [27]. For both the ROC and PR cases, the extensions are equivalent to randomly guessing edge existence for the missing part of the predicted network, which is trivial to perform, and thus guaranteed to be possible to realise. GENIE3 never infers self regulatory interactions for genes. To avoid excessively penalizing this, the benchmarking of GENIE3 predictions in this publication excludes self regulatory interactions. For all other methods, they are included. All parameters used for all aspects of the method performance evaluation are included with the publication reproduction code.

### Processing of ENCODE reference data

shRNA-seq perturbation experiments from the ENCODE project were used as reference for properties of simulated data. Perturbation data from the K562 and HepG2 cell lines and corresponding unperturbed control samples were downloaded from ENCODE [28]. Gene expression in terms of transcript abundance was quantified using pseudo alignment with Salmon [29]. To calculate log2 fold-change (log2FC) values, perturbation-induced expression values were compared to the controls, and technical replicates were handled by averaging. Where multiple perturbations targeted the same gene, they were averaged to a single log2FC value. Code and detailed description of the data processing is available in the publication reproduction code repository (Code Availability section).

## Results

GeneSNAKE is a simulation tool that relies on an ODE model to capture the dynamic behavior of gene regulatory interactions. Building on previous work, GeneSNAKE’s ODE system shares many features with other simulators. Although this methodology has often been used to simulate data for GRN inference, only limited testing has been done to assess how well it models the dynamics of a real system. To address this and demonstrate that GeneSNAKE is capable of capturing the complex dynamics observed in biological experiments, we tested how well the simulated data correlates with experimental data derived from work by Hackett et al. [30]. This experiment studied the effect of overexpressing selected transcription factors in *S. cerevisiae*. We tested whether GeneSNAKE can capture the same effect by simulating the same experiments and measuring how well the simulated data correlates with the experimental data. The test was performed on a sub-selected set of genes, measuring the effect in only those genes that were directly regulated by the perturbed transcription factor or those regulated by a directly targeted gene. This was to ensure that the expression of genes in the system changed so as to avoid issues with a false low or high correlation in genes with no detectable change. The test revealed that in a majority of the genes (146 of 250 tested) GeneSNAKE was able to perfectly capture (Spearman correlation 1.0) the dynamics of their behavior in biological data. On average the GeneSNAKE simulation had a Spearman correlation of 0.62 indicating that the simulation largely follows the trends seen in experimental data. **Figure 2** shows the distribution of correlations for each subset GRN that was investigated. It also shows the dynamics of one GRN in detail, which consists of the AFT2 transcription factor and its downstream genes. It can be seen that while the absolute expression levels are generally not the same, the behavioral trends are similar.

**Figure 2.**
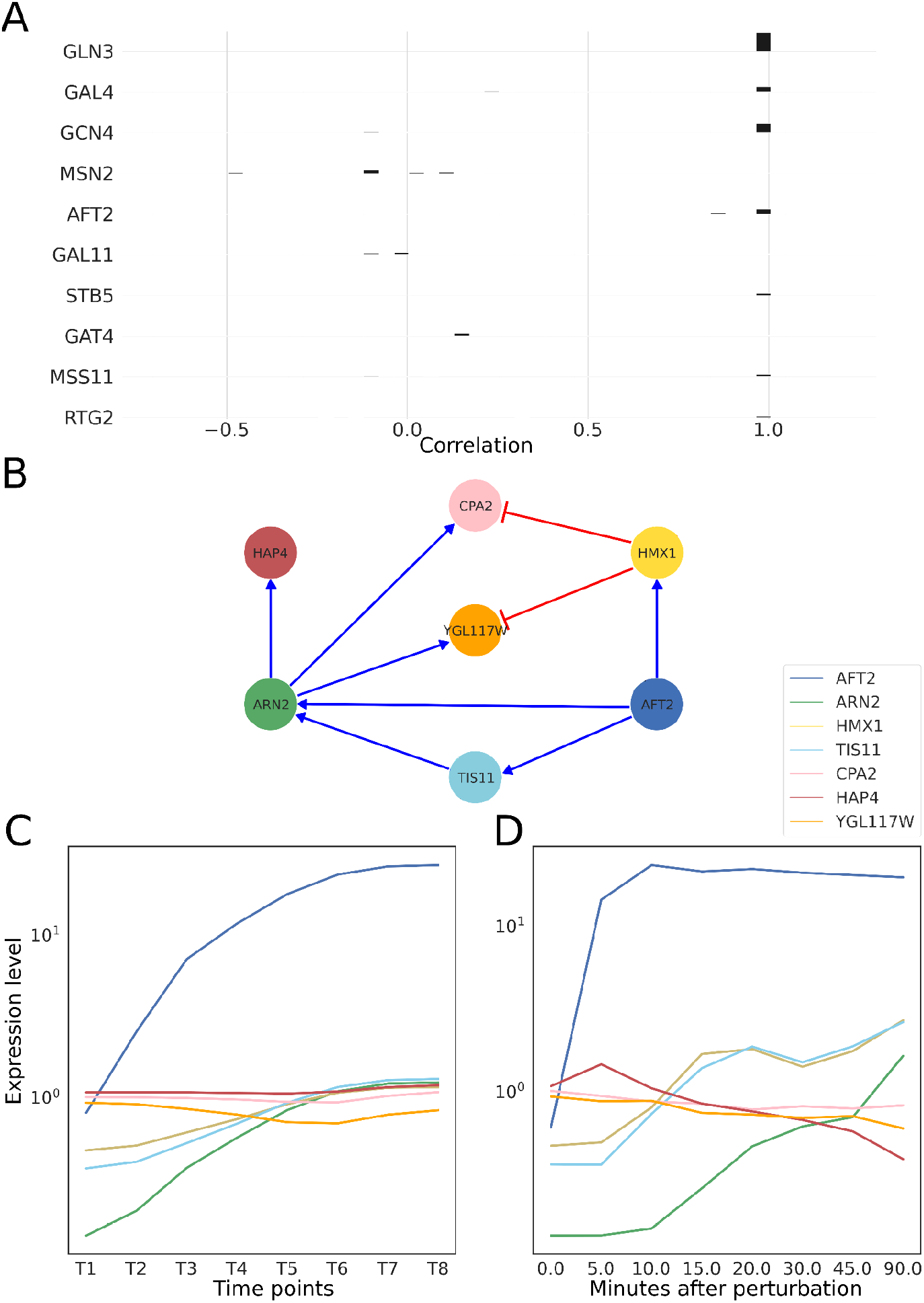
Correlation between GeneSNAKE simulation and experimental data. Subsets of the GRN model from Hackett et al. were used to simulate the system’s evolution over time. (A) The left column lists the regulators that were perturbed by overexpression and the blue bars display a histogram of correlations between measured and simulated expression of downstream genes for each perturbed regulator. The image shows an overall good correspondence over time between gene expression in the two datasets indicating that GeneSNAKE can model biologically relevant data. (B) An example subset GRN for the regulator AFT2 and downstream genes. (C) Expression levels simulated over time by GeneSNAKE for the AFT2 subset GRN and the Hackett et al. ODE model. Note that the simulation goes to a steady-state. (D) Expression levels measured experimentally in yeast over time by Hackett et al. for the AFT2 subset GRN genes. Note that their time series, unlike the simulation in C, does not go to a steady-state, hence should be compared to the first part of the simulation. Also, note that overexpression of AFT2 in the Hackett et al. experimental data is faster than in the simulated data due to their use of an artificial induction system for rapid overexpression.

Next, we wanted to demonstrate that GeneSNAKE can simulate data with a meaningful connection between the underlying GRN and the simulated data. To test this, GRNs were created using the GeneSNAKE GRN generation tool with the FFLatt algorithm. The GRNs were generated with negative self-loops to ensure system stability. GeneSNAKE was then used to generate an ODE model and simulate single perturbation steady-state data from this network with one perturbation for each gene across three replicates. A range of Signal to Noise Ratio (SNR) levels were applied, going from very high to very low amounts of added noise, where SNR is defined as:

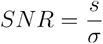

where *S* is the expression of a given gene in a given experiment which is multiplied with noise sampled from a Gaussian with a mean of 1 and a standard deviation of *σ*. **Figure 3** shows that it is possible to infer accurate networks from GeneSNAKE data, with a clear trend of increasing correctness of the GRN inference methods as the SNR increased. The reason that AUROC does not reach 1 even with no added noise can be attributed to the fact that GeneSNAKE’s data generation uses a non-linear model with stochastic components, while the inference methods use linear models. Furthermore, we generated the GRNs with FFLatt that induces a network topology with many feed-forward motifs, which makes the inference task harder.

**Figure 3.**
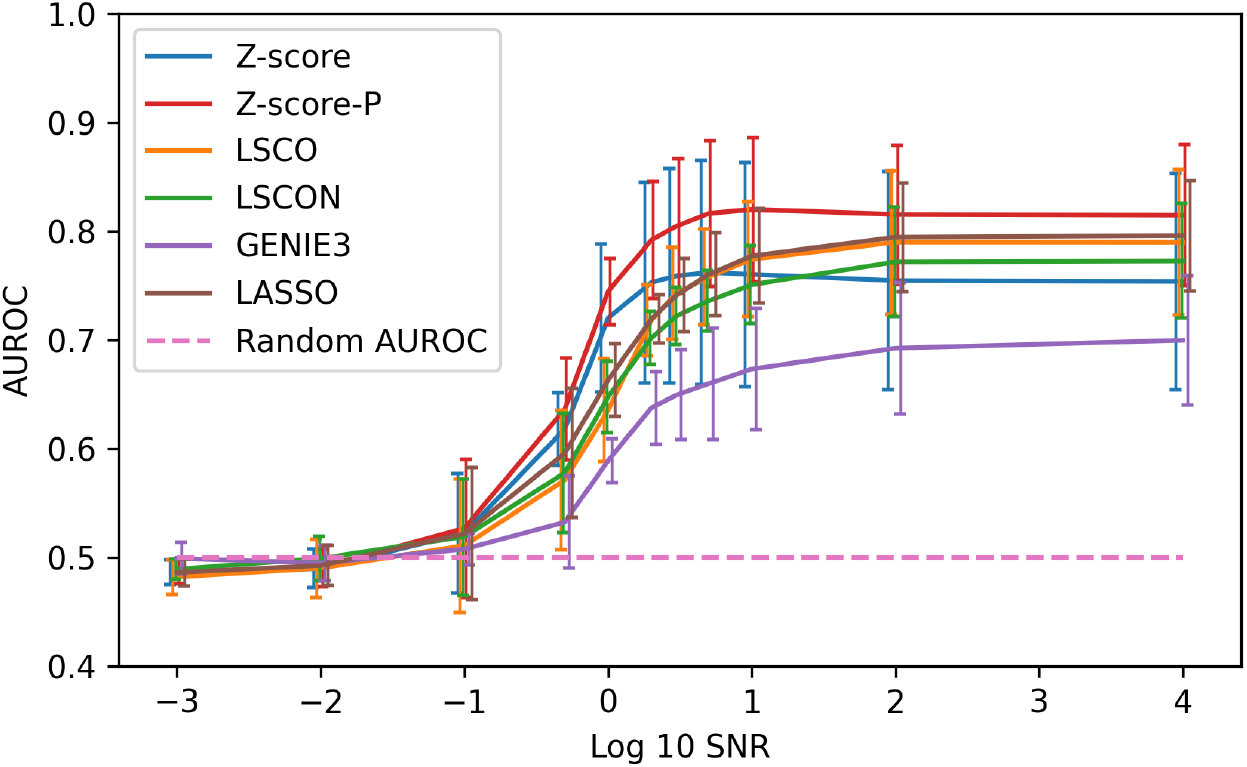
GRN inference accuracy for six inference methods using data produced by GeneSNAKE. Steady-state data was simulated with GeneSNAKE from FFLatt GRNs containing 100 genes, for eight different Signal to Noise Ratio (SNR) levels. Accuracy was measured in terms of Area Under Receiver Operating Characteristic (AUROC).

To test how well a GeneSNAKE simulation corresponds to real data, perturbation-based data for cell lines K562 and HepG2 from ENCODE was used [28]. Fold-change expression matrices were computed, where each gene is knocked down in two replicate experiments, for a total of 232 genes and 464 experiments (Figure S1, methods, and publication reproduction code repository). A network with the same number of genes was simulated using the FFLATT algorithm, and expression data was generated using the same perturbation design. A comparison of simulated and real data was made using the exploratory data analysis (EDA) tool provided in GeneSNAKE, giving an overview of general data set properties and sample statistics as well as visualizations of the above. **Figure 4** shows that GeneSNAKE is able to generate GRNs and with them simulate data which has properties similar to real gene expression data.

**Figure 4.**
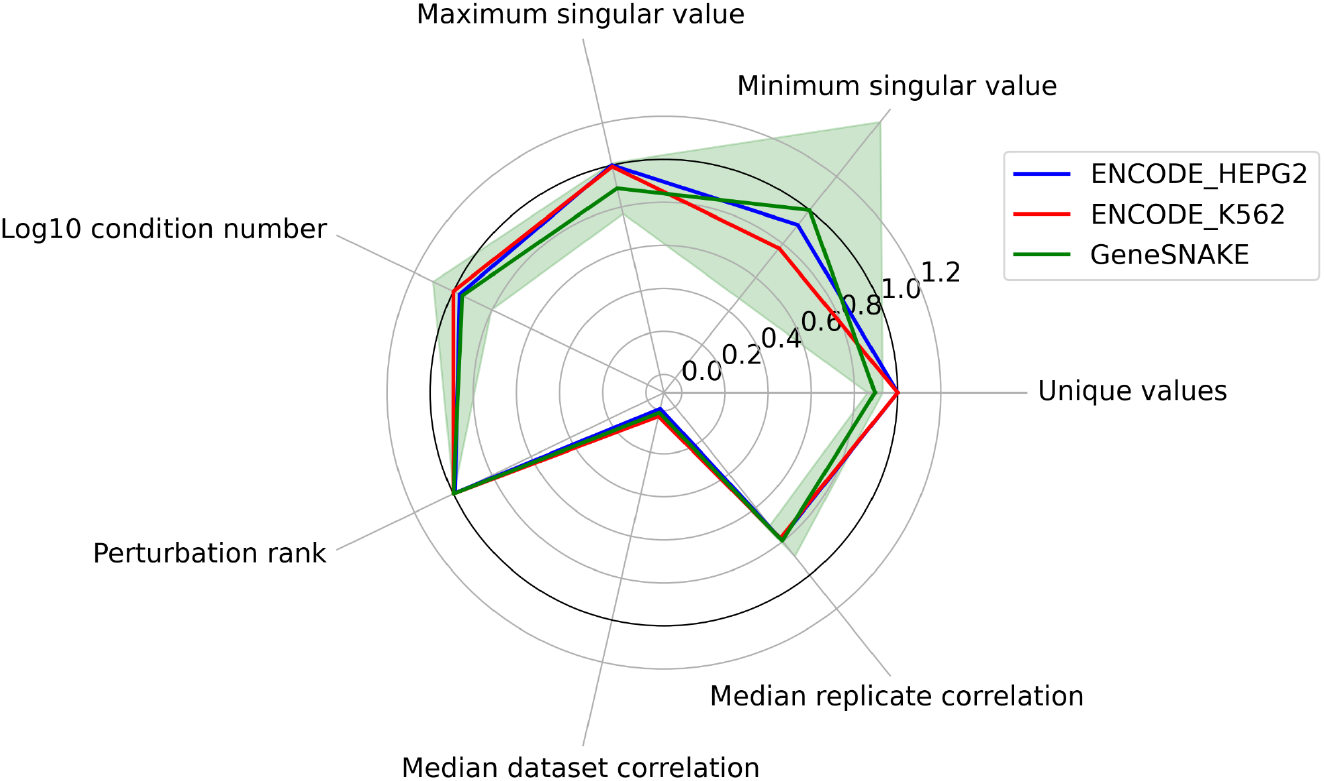
GeneSNAKE can generate realistic perturbation-induced gene expression data. Comparison of data properties from real (ENCODE cell lines K562 and HepG2) and GeneSNAKE simulated data of the same size. All datasets contain log fold changes at steady state after single gene knockdowns. Fraction unique values, median replicate correlation, median data set correlation, and perturbation rank are shown as original values, while log 10 condition number and singular values are shown as the proportion of the respective maximum value among the three datasets. Perturbation rank indicates the specificity of the knock-down to the target gene. A rank of 1 means that the target was most knocked down of all genes. Shown is the median fractional knock-down rank across all genes.

Another key feature of a simulation tool is scalability, which we tested by evaluating the run time for increasing GRN size. **Figure 5** shows that the GeneSNAKE data generator scales about *O*(n^3^) with the size of the system, for instance simulating data with 500 genes takes about 50 minutes while simulating data from 1000 genes takes about 3 hours. Furthermore, the test shows that a majority of this time is spent on solving the ODE system once the GRN has been generated. The simulation was run on an 8-processor CPU (Intel i7-6700).

**Figure 5.**
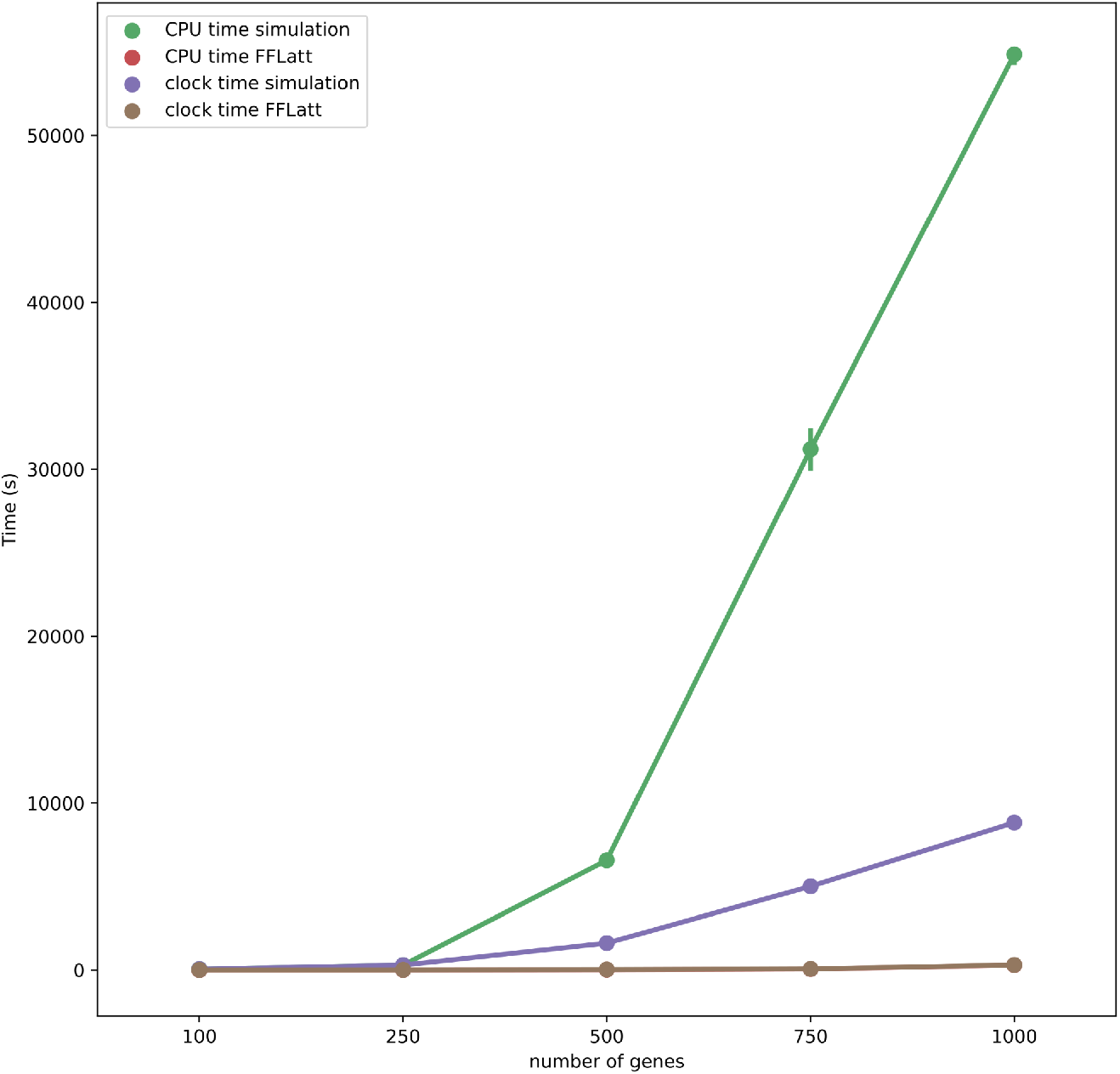
Run time in seconds for GeneSNAKE simulations over varying GRN sizes. The difference between CPU time and wall clock time shows that GeneSNAKE, by relying on parallelization, can drastically lower the real-time of the simulation. Furthermore, it shows that solving the ODE system stands for almost all of the runtime. The reason for this is that the ODE model by its nature must be solved sequentially, giving it a high time complexity. The CPU time for FFLatt is the same as the clock time. The simulation was run on an 8-processor CPU (Intel i7).

## Discussion

We here present GeneSNAKE, a new Python-based package for generating perturbation-induced gene expression data for a given gene regulatory network. The tool is able to generate both GRNs and expression data, as well as perform benchmarking of predicted GRNs compared to ground truth GRNs. We show that the GeneSNAKE ODE model is capable of capturing the same trends over time that are observed in experimental data using the same underlying network, as well as generating data with properties similar to biological data. We also show that the generated data is related to the original GRN and that the noise models can add meaningful noise, allowing developers to explore how different data qualities affect their inference method. GeneSNAKE’s main novelties are the customizable perturbation model, a wide range of perturbation schemes, control of the noise level, and several noise models, in an open-source Python package.

We used GeneSNAKE to simulate data given a known GRN, and compared the time-dependent dynamics of the simulation with an experimental time series of gene expression. It was observed that GeneSNAKE data correlates well with experimental data, suggesting that the main trends of the experiment are accurately represented when generating data from known biological GRNs. A finding that, alongside the flexible perturbation solution and noise models, highlights GeneSNAKE’s usefulness in not only generating data from simulated GRNs for benchmarking purposes but also to explore the behaviour of experimental systems.

When testing for the ability to generate benchmarking data a clear trend was observed that the performance of all methods increases with decreasing noise, an expected finding that has been previously shown [31]. A more unexpected finding was that those methods that rely on simpler algorithms, especially the Z-score method which uses Z-score transformation of the data to select the most relevant edges, outperformed methods with more complex calculations. The same trend has been observed before [31], in particular the trend of more complex methods underperforming with very noisy data.

Compared to other simulation packages such as GeneNetWeaver, an advantage of GeneSNAKE is that it gives much better control over GRN and data properties, and experimental design. Properties such as sparsity, stability, scale-freeness, condition number, variance, and noise level can have a different impact on different inference methods and therefore need to be varied in a controlled fashion. The GeneSPIDER package allows control of many of these properties but, like many other simulators and inference methods, simplifies the dynamics of gene regulation by only considering the steady-state, when the system no longer changes. Another issue with GeneNetWeaver is that it by default generates GRNs with link weights of 1, 0, or -1, which limits the quantitativeness of the system. With GeneSNAKE we combine the high controllability of GRN and data properties with time-dependent dynamical modeling as in GeneNetWeaver to obtain realistic yet flexible modeling. GeneSNAKE further has more advanced control of regulator collaboration, perturbation strength, and perturbation schemes.

Another important strength of the GeneSNAKE tool is the addition of FFL motifs in the network generation. These motifs have been shown to be enriched in biological GRNs [12,32,33] and have in previous benchmarks been shown to affect the performance of various models in different ways. For example, the large community-driven benchmark DREAM 5 noted that even the best-performing regression models struggled to identify motif structures while information theory models could to a greater extent recover these motifs [3]. This hints at the importance of using realistic GRNs for simulation, as unrealistic GRN properties could lead to invalid performance comparisons of GRN inference methods, especially if they have different abilities to cope with motif biases.

## Key points

- GeneSNAKE is a Python package for generating biologically realistic GRNs and perturbation-induced expression data, mainly for benchmarking.
- Synthetic expression data follows the dynamics specified in an ODE model based on the GRN and a perturbation design.
- It offers extensive control over network and data properties, with support for diverse noise models, a wide range of perturbation schemes, and tunable levels of noise and perturbation strength.
- The package includes benchmarking functions to compare networks and analyze properties of data and GRNs, helping create more realistic simulated datasets.

## Acknowledgments

This work was supported by the Swedish Research Council grant 2019-04095.

## Author contributions

T.H. implemented a major part of the tool, performed analysis, and constructed figures 1 and 5. A.B. performed analysis, implemented functions of the tool, and constructed figures 3 and 4. N.L. performed analysis, implemented functions of the tool, and constructed figure 2. E.K.Z. implemented functions of the tool, performed analyses, and processed ENCODE data for figure 4, M.G. implemented functions of the tool and created the GeneSNAKE logo. E.S. conceived and supervised the work. T.H., A.B., N.L., E.K.Z, M.G., and E.S. wrote the manuscript. All authors reviewed and approved the final version of the manuscript.

## Code availability

GeneSNAKE is publicly available as a Python package at: https://bitbucket.org/sonnhammergrni/genesnake/

Code for reproducing the figures and data processing in this publication is available at: https://bitbucket.org/sonnhammergrni/genesnake_publication_reproduction/

## Data availability

The ENCODE data for HepG2 and K562 cell lines are publicly available via the ENCODE [28] project website https://www.encodeproject.org/. See the publication reproduction code repository for a detailed description of its accession and processing.

The data for the work by Hackett et al., 2020 is available at the IDEA project website: https://idea.research.calicolabs.com/data

## Notes

https://idea.research.calicolabs.com/data

https://www.encodeproject.org/

### Competing Interest Statement

The authors have declared no competing interest.

