## Supplemental material for "GeneSNAKE: a Python package for benchmarking and simulation of gene regulatory networks and perturbation-induced expression data"

### Supplementary text

#### Data summary

To investigate perturbed data properties, we created a set of visualizations that characterize density, normality, values distributions, and perturbation ranking. First, data density is shown as a distribution of values via kernel density estimate. Data normality is shown as a Q-Q plot where the P value from the Kolmogorov–Smirnov test is displayed. Next, the distribution of data values is given as a barplot where bars represent fractions of zero, infinite, NaN, positive, and negative values. Finally, perturbation ranking displays a histogram of perturbation fractions that represent the percentage of genes having lower expression than the perturbed gene for each experiment. In other words, perturbation ranking characterizes how well the perturbations were performed in experiments.

#### Benchmarking

GeneSNAKE allows benchmarking of an inferred set of networks against a gold standard network. By comparing adjacency matrices it calculates the confusion matrix where true positives (TP), true negatives (TN), false positives (FP), and false negatives (FN) are summed up according to the presence of corresponding links. Afterward, the confusion matrix is used to estimate various performance measurements (Table S1) such as Area Under the Receiver Operating Characteristics (AUROC), Area Under the Precision-Recall (AUPR), sensitivity (TPR), specificity (TNR), precision (PPV), negative predictive value (NPV), miss rate (FNR), fall-out (FPR), false discovery rate (FDR), false omission rate (FOR), positive and negative likelihood ratio (LRP, LRN), prevalence threshold (PT), threat score (TS), F1-score, Matthew's correlation coefficient (MCC), Fowlkes-Mallows index (FM), informedness (BM), markedness (MK), and diagnostic odds ratio (DOR).

**Table S1.** Performance measurements implemented in GeneSNAKE for network benchmarking.

| measurement | | | | | | | | |
|---|---|---|---|---|---|---|---|---|
| TPR | TNR | PPV | NPV | FNR | FPR | FDR | FOR | LRP |
| $\frac{TP}{TP+FN}$ | $\frac{TN}{TN+FP}$ | $\frac{TP}{TP+FP}$ | $\frac{TN}{TN+FN}$ | $\frac{FN}{FN+TP}$ | $\frac{FP}{FP+TN}$ | $\frac{FP}{FP+TP}$ | $\frac{FN}{FN+TN}$ | $\frac{TPR}{FPR}$ |
| measurement | | | | | | | | |
| LRN | PT | TS | F1-score | MCC | FM | BM | MK | DOR |
| $\frac{FNR}{TNR}$ | $\frac{\sqrt{FPR}}{\sqrt{TPR}+\sqrt{FPR}}$ | $\frac{TP}{TP+FN+FP}$ | $\frac{2*TP}{2*TP+FP+FN}$ | $\sqrt{PPV * TPR * TNR * NPV} - \sqrt{FDR * FNR * FPR * FOR}$ | $\sqrt{PPV * TPR}$ | $TPR + TNR - 1$ | $PPV + NPV - 1$ | $\frac{LRP}{LRN}$ |

#### Sequencing noise model

Inaccurately performed sequencing experiments may introduce a bias in gene expression measures (Ross et al. 2013). It may be caused by poorly designed experiments, GC-rich sequences, or various technical errors that lead to unmappable reads or over-mapping certain regions with reads. Furthermore, gene length is a factor that may affect the sequencing coverage. It has been concluded that short gene length bias was detected for scRNA-seq (Phipson, Zappia, and Oshlack 2017) similarly to bulk RNA-seq (Oshlack and Wakefield 2009).

To model synthetic data that follow the real data properties, we developed a coverage-based noise model. To achieve this, we imitate a sequencing process relying on the Lander-Waterman equation that allows us to estimate the sequencing coverage $C$ as follows: $C = \frac{L \times N}{G}$, where $L$ is the read length, $N$ is the total number of reads, and $G$ is the total length of a sequenced DNA. In this work, the values of $C$ and $L$ are assumed based on literature to set it according to a given sequencing technique, e.g. average coverage of 5 for RNA-seq studies (McIntyre et al. 2011). Moreover, sequencing error rates and gene lengths are also taken from literature or public databases. Some other parameters, such as the number of reads can be estimated based on the Lander-Waterman equation: $N = \frac{C \times G}{L}$.

To model the mapping procedure, we randomize the positions of aligned reads from a uniform distribution. Here, we take real gene start and end positions based on the human genome to establish their length. Afterward, we assume that the count value for a single gene in a given experiment is equal to the number of reads that cover the gene (Pepke, Wold, and Mortazavi 2009). This is referred to as simulated counts. Next, to simulate multiple experiments (or samples) this procedure is repeated a certain number of times. In addition, the entire process can be again repeated to simulate technical replicates. However, the described procedure simulates mainly subtractive noise and just a small fraction of additive noise from a GC-content-related step. Thus, the next stage of noise modeling involves the negative binomial (NB) distribution to simulate the distribution of counts as additive noise. Furthermore, zero inflation is introduced to data to simulate dropout rates. It is together known as a zero-inflated negative binomial (ZINB) distribution used in noise modeling for single-cell RNA-seq data (Eraslan et al. 2019). Finally, a matrix of error ratios is obtained by element-wise matrix operations as follows:

$$E_{seq} = ((counts_{simulated} + E_{NB}) \oslash counts_{expected}) \odot E_{ZI} \quad \text{(Eq. S1)}$$

where $counts_{simulated}$ is a matrix of simulated counts, $E_{NB}$ is a matrix of erroneous counts drawn from the negative binomial distribution, $counts_{expected}$ is a matrix of theoretical counts estimated based on the Lander-Waterman equation for each gene, and $E_{ZI}$ is a binary matrix for zero inflation. To obtain a noise model with dropouts mimicking single-cell data, set the probability for $E_{ZI}$ greater than 0. Otherwise, set it to 0 to obtain dropout-free data mimicking bulk data. The error matrix is then used in a multiplicative way on noise-free data.

#### Supplementary figures

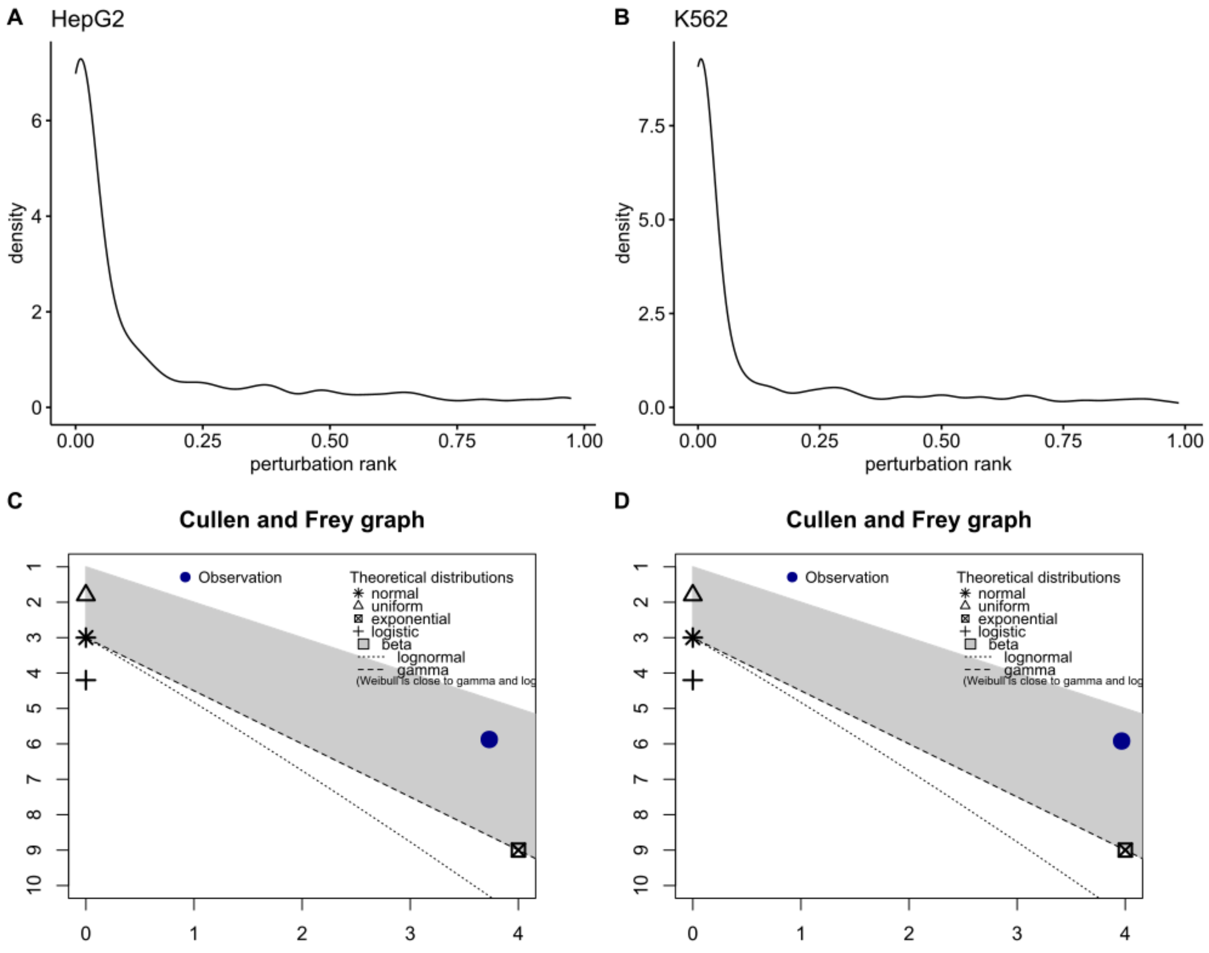

**Figure S1.** Perturbation rank density and distribution for **A.** HepG2 and **B.** K562 cell lines. The type of distribution for perturbation rank was estimated with the Cullen and Frey graph for **C.** HepG2 and **D.** K562 cell lines. Observations indicate beta distribution.

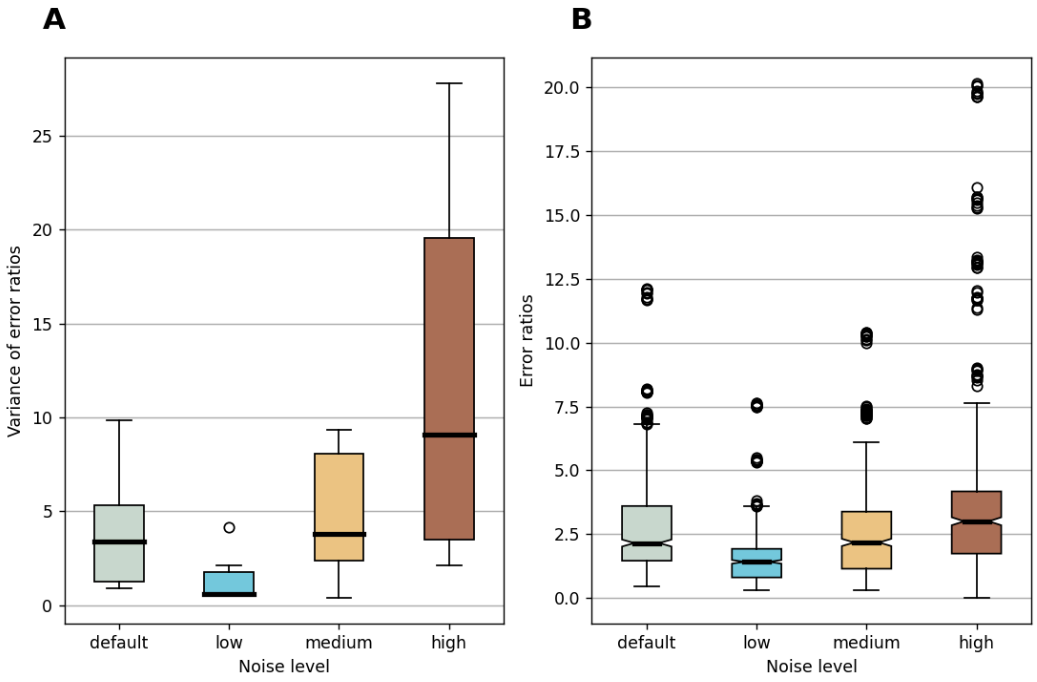

**Figure S2.** Performance of various noise options in sequencing-like noise by changing maximum alignment error (ermax), negative binomial probability (prob_nb), and variance scaling factor (sf). Some of the parameters were left default: minimum alignment error (ermin), technical replicates number (techrep), and probability of zero-inflation (prob_zi). The following settings were used: default (ermin=0.45, ermax=0.73, prob_zi=0, prob_nb=0.5, sf=0.75), low (ermax=0.6, prob_nb=0.6, sf=0.8), medium (ermax=0.7, prob_nb=0.5, sf=0.6) and high (ermax=0.8, prob_nb=0.4, sf=0.4) **A.** Variance of the error ratio matrix $E_{seq}$ and **B.** General statistics of the error ratio matrix $E_{seq}$. As ratios are randomly generated, each run was repeated six times. This analysis was performed for 10 randomly selected genes for a randomly generated count matrix of size 10x10.

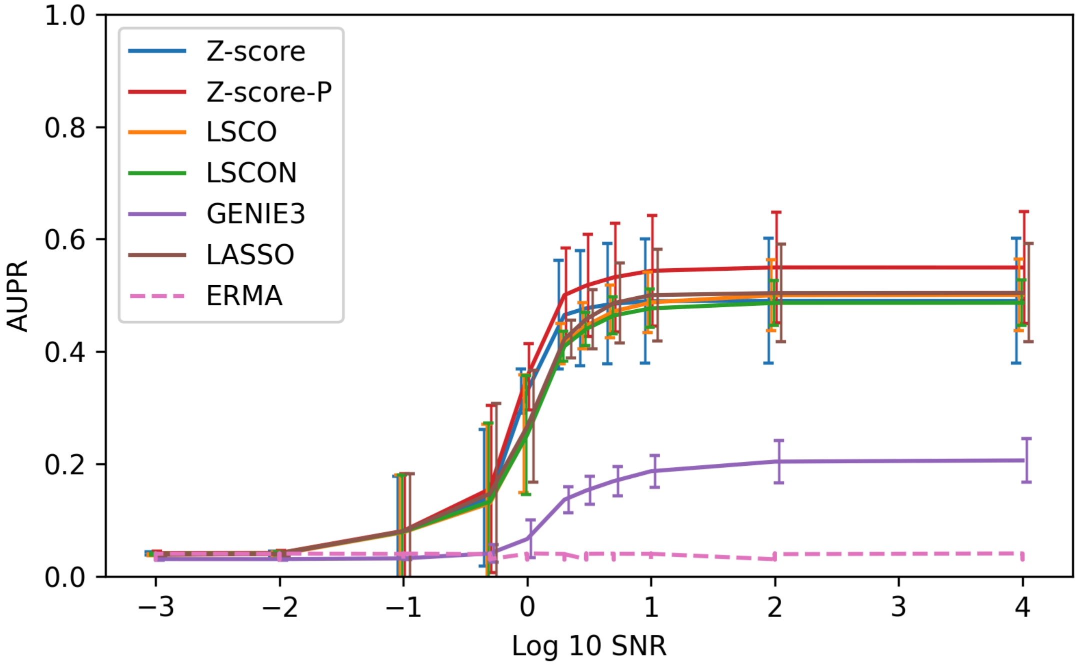

**Figure S3.** GRN inference accuracy for six inference methods using data produced by GeneSNAKE. Steady-state data was simulated with GeneSNAKE from FFLatt GRNs containing 100 genes, for eight different Signal to Noise Ratio (SNR) levels. Accuracy was measured in terms of Area Under Precision Recall (AUPR). Error bars are ± 1 standard deviation based on 7 different networks. The error bars are slightly offset along the x axis for visual clarity. The same SNR levels were recorded for all model types.

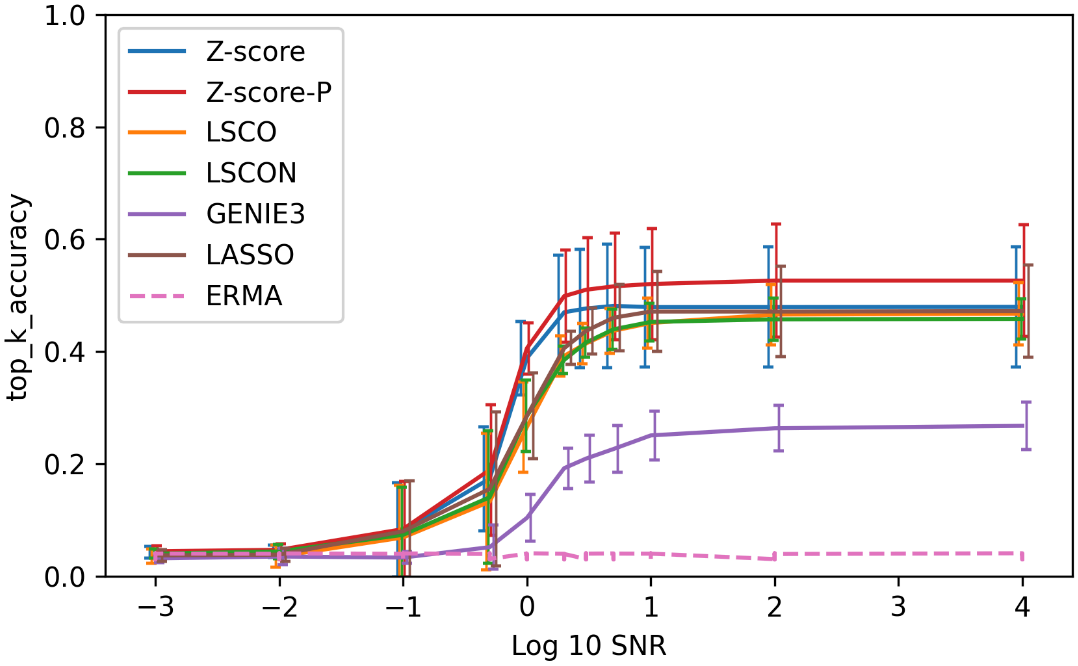

**Figure S4.** GRN inference accuracy for six inference methods using data produced by GeneSNAKE. Steady-state data was simulated with GeneSNAKE from FFLatt GRNs containing 100 genes, for eight different Signal to Noise Ratio (SNR) levels. Accuracy was measured in terms of top K accuracy, the accuracy of the top K most confidently predicted links, with K equalling the number of existing edges in the reference network. Error bars are ± 1 standard deviation based on 7 different networks. The error bars are slightly offset along the x axis for visual clarity. The same SNR levels were recorded for all model types.

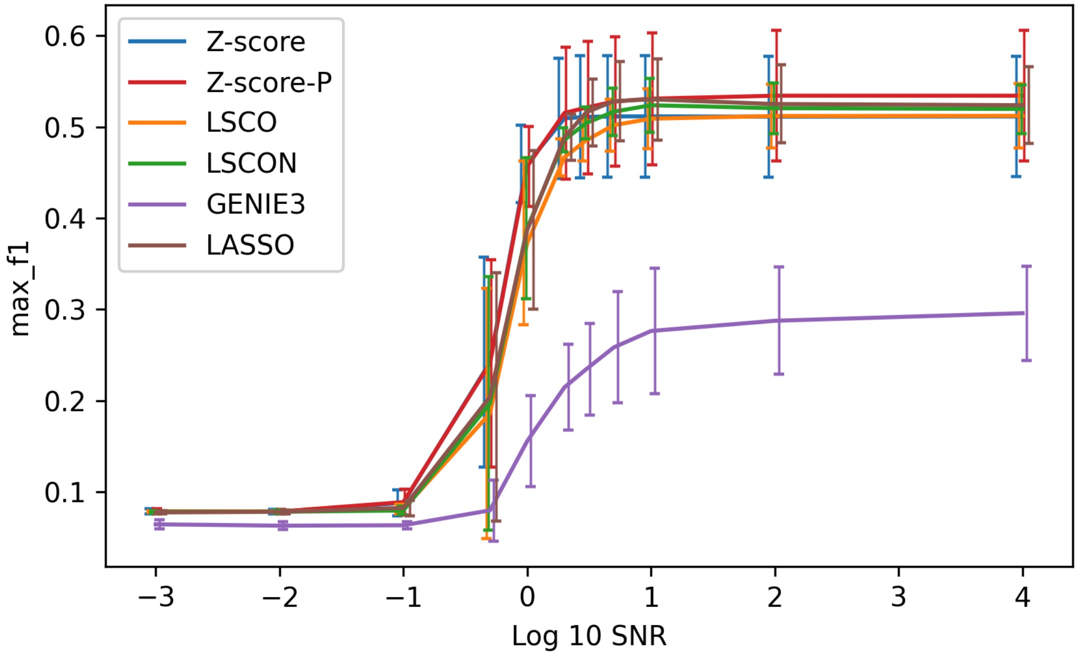

**Figure S5.** GRN inference accuracy for six inference methods using data produced by GeneSNAKE. Steady-state data was simulated with GeneSNAKE from FFLatt GRNs containing 100 genes, for eight different Signal to Noise Ratio (SNR) levels. Accuracy was measured in terms of maximum F1 score over thresholds for all values in the estimated networks. Error bars are ± 1 standard deviation based on 7 different networks. The error bars are slightly offset along the x axis for visual clarity. The same SNR levels were recorded for all model types.

McIntyre, Lauren M., Kenneth K. Lopiano, Alison M. Morse, Victor Amin, Ann L. Oberg, Linda J. Young, and Sergey V. Nuzhdin. 2011. “RNA-Seq: Technical Variability and Sampling.” *BMC Genomics* 12 (June): 293.

Oshlack, Alicia, and Matthew J. Wakefield. 2009. “Transcript Length Bias in RNA-Seq Data Confounds Systems Biology.” *Biology Direct* 4 (April): 14.

Pepke, Shirley, Barbara Wold, and Ali Mortazavi. 2009. “Computation for ChIP-Seq and RNA-Seq Studies.” *Nature Methods* 6 (11 Suppl): S22–32.

Phipson, Belinda, Luke Zappia, and Alicia Oshlack. 2017. “Gene Length and Detection Bias in Single Cell RNA Sequencing Protocols.” *F1000Research* 6 (April): 595.

Ross, Michael G., Carsten Russ, Maura Costello, Andrew Hollinger, Niall J. Lennon, Ryan Hegarty, Chad Nusbaum, and David B. Jaffe. 2013. “Characterizing and Measuring Bias in Sequence Data.” *Genome Biology* 14 (5): R51.
